# Geometrical factors determining dendritic domain intersection between neurons: a modeling study

**DOI:** 10.1101/2022.03.14.484202

**Authors:** Rafael Ignacio Gatica, Trinidad Montero, Navid Farassat, Pablo Henny

## Abstract

Overlap between dendritic trees of neighboring neurons is a feature of nervous systems. Overlap allows neurons to share common afferences and defines the topography of the circuit they belong to. Proximity is also a requirement for dendritic communication, including dendro-dendritic synaptic contacts. We simulated overlap dynamics between pairs of ventral tegmental area dopamine neurons, a population characterized by diverse and extensive dendritic domains. Using each neuron’s 3D convex hull (CH) as a proxy for dendritic domain size and shape, we examined intersection versus cell-to-cell distance curves for 210 pairs of neurons, and found that decay dynamics were diverse and complex, indicating that intersection between real dendritic domains does not comply with spherical or isotropic shape assumptions. We re-examined intersection dynamics for the same 210 pairs, but this time with either: a) a normalized volume corresponding to the average volume of all CHs, b) normalized shapes, using an average CH shape, or CHs from neurons exhibiting average dendritic distribution, isotropy or morphology, or c) a normalized cell body position, which we artificially placed in each CHs’ centroid. All three interventions significantly increased pair intersection and simplified decay dynamics, yet shape normalization had the strongest influence. Critically, shape uniformity was also the most relevant factor for increased dendritic domain pair intersection using the neurons’ original brain position and distances. We applied this experimental approach to other populations using reconstructions from neuromorpho.org database and found that shape and intersection dynamics were cell-type dependent. We conclude that intersection between neurons is not only maximized by proximity, but that individual-specific dendritic domain geometries have a profound impact too. The results predict that one biological solution for circuits requiring selective connectivity is to exhibit greater heterogeneity in size, cell body location, and especially shape, among their constituent neuronal elements.

## 1. INTRODUCTION

Overlap between the dendritic trees of adjacent neurons is a common feature of the nervous system (Harris & Spacek, 2017; Nieuwenhuys et al., 1998; Ramon y Cajal, 1904). Neurons whose respective dendrites share a given locus are likely to share common afferent inputs (Braitenberg & Schüz, 1998; Packer et al., 2013; Peters & Feldman, 1976). Also, the topographical organization of a projection system is partly defined by the degree to which spatially adjacent neurons share or exclude sensory or afferent inputs (Nieuwenhuys et al., 1998). The role of dendritic fields’ overlap may be particularly important in brain structures such as the neocortex of large mammals, which in large-brain exhibit a trend to decreased neuronal density (neuronal cell body density) and a more extensive neuropil (Herculano-Houzel, 2011). In addition, proximity also ensures dendro-dendritic paracrine transmission (Brombas et al., 2017; Rice & Patel, 2015) and it is a requirement for establishing dendro-dendritic synapses (Groves & Linder, 1983; Hinds, 1970). The role of dendritic overlap between adjacent neurons in determining integration or segregation of afferent signals may play is also expected to be critical in species in which neuronal density (neuronal cell bodies) decreases and neuropil increases, as it happens in the neocortex of large-brain mammals (Herculano-Houzel, 2011)

Dendritic domains overlap has been studied at a population level in somatosensory and visual systems in the context of neurons’ receptive field development (Jan & Jan, 2010; Lefebvre et al., 2015; Oleson et al., 2014). In invertebrates, for instance, initial overlap between same-class dendritic arborization skin surface neurons is gradually eliminated, resulting in a complete and non-overlapping coverage of the sensory surface, and similar processes have been described for ganglion and some amacrine cell types in the vertebrate retina (Jan & Jan, 2010; Lefebvre et al., 2012, 2015). Conversely, neurons whose function requires signal integration from multiple receptive fields, as it is the case for motion detection cholinergic or dopaminergic amacrine cells, exhibit dendritic domains that overlap extensively (Brombas et al., 2017; Farajian et al., 2004; Keeley & Reese, 2010).

As for other central nervous system neurons, only a few studies have expressly examined and reported dendritic domain overlap and its anatomical or developmental determinants. For instance, overlap between mouse Purkinje cells was shown to depend on lobule location and developmental stage (Nedelescu et al., 2018) and, in mouse cortical pyramidal cells, overlap was shown to be severw and dendritic arborization development (Zhang et al., 2012)

In this study we sought to understand overlap in central nervous system neurons using a single cell pair analysis approach, with which we aimed to capture the fine dynamics that underlie intersection in a real, three-dimensionally organized and morphologically heterogeneous population. More specifically, we sought to answer the following questions: 1) what are the geometrical determinants of dendritic domain intersection between pairs of neurons in adult central neurons? 2) what is the specific influence of dendritic domain size, shape and the cell body placement on intersection? 3) how does cell-to-cell morphology heterogeneity affect dendritic domain pair intersection dynamics?

To address these questions, we studied adult mouse ventral tegmental area (VTA) dopamine (DA) neurons, a population involved in reinforcing behavior, and whose neurons are morphologically characterized by long radiating and overlapping dendrites (Montero et al., 2021) that sub-serve integration of multiple and heterogeneous input sources (Geisler & Zahm, 2005). We used the neurons’ convex hulls polyhedrons (CHPs) generated from previously 3D reconstructed VTA DA neurons (Montero et al., 2021). CHPs enclose the minimal convex volume in which the dendritic domains locate and have been used as proxies for dendritic domain size and shape (Felix et al., 2013; Gärtner et al., 2004; Gertler et al., 2008; Ito, 2020; Malmierca et al., 1995; Vrieler et al., 2019).

Consequently, in this study we set the following aims: 1) to test the assumption that CHPs are adequate proxies for dendritic domain size and shape and, similarly, that convex hulls intersections are adequate proxies for dendritic domain and proximity, 2) to model the influence of cell body to cell body distance on intersected volume in the mediolateral (ML), dorsoventral (DV) and anteroposterior (AP) axes, 3) to model the influence of CHPs size, shape and cell body placement inside the dendritic domain, on intersection, 4) to test whether results are replicable when examining natural pair intersection using the original placement of neurons inside the brain and, finally, 5) to test whether these methodologies are suitable to study intersection dynamics in other neuronal populations.

As shown below, we provide evidence that supports the convex hull assumption for dendritic shape and intersection and describe that pair to pair intersection dynamics are diverse and complex. We then go on to demonstrate a particularly important role of neurons CHP shape homogeneity in dendritic fields intersection. Finally, we report that this experimental approach can be used in other central neuronal populations exhibiting diverse sizes and morphologies, and that, regarding VTA DA neurons, intersection principles revealed by modeling are replicated when studying neurons in their original brain location.

## 2. MATERIAL AND METHODS

### 2.1 Reconstructed neurons

Reconstructed DA neurons came from a pool of previously reported mouse neurons (VTA, n = 15; SNc, n = 15, (Farassat et al., 2019; Meza et al., 2018; Montero et al., 2021)). Briefly, neurons were labeled with Neurobiotin using juxtacellular or intracellular labeling methods and brain tissue was processed to reveal Neurobiotin and tyrosine hydroxylase to confirm the dopaminergic profile. Next, neurons were manually reconstructed using Neurolucida (MBF Bioscience) from confocal image stacks (more details in (Montero et al., 2021)). The somatodendritic domain was completely filled, shrinkage corrected and reconstruction tilted-corrected. For some analyses (Figure 2B, Figure 8), individual neuronal reconstructions were placed in a custom-made 3D map of the VTA based on (Fu et al., 2012), according to the ML, DV, and AP location of the cell body (Figure 8A).

Also, neurons from other studies were used from the neuromorpho.org database, which fulfilled the following criteria: a) reconstructed from in vivo experiments such that no dendrites are lost, b) somatodendritic domains were completely filled, c) shrinkage correction was applied and d) orientation of neurons in relation to the cartesian space was known. The following neurons were used: SNc DA neurons (rat, n = 10, (Henny et al., 2012)), cerebellum, anterior lobule V of vermis (PC, mouse, n = 10, (Nedelescu et al., 2018)), mediodorsal thalamic nucleus neurons (MD, rat, n = 5, (Kuramoto et al., 2017)) and spinal motoneurons (MN, rat, n = 3 (Rotterman et al., 2014)).

### 2.2 Morphological analyses

Morphological and 3D wedge data were extracted using Neurolucida Explorer software (MBF Bioscience). For wedge analysis, the distribution of the dendritic processes as they extend from the cell body in every direction was obtained. To achieve this, the neuron is placed in the center of a cylinder, which in this study was divided into 4 angular wedges, and in 2 parallel planes perpendicular to the direction of the cylinder, with the division of the planes centered in the cell body. For instance, if the cylinder is placed parallel to the anteroposterior axis, the four angular wedges report the location of the processes in the dorsomedial, ventromedial, ventrolateral, and dorsolateral quadrants, at each of the 2 planes defined in the AP axis (Figure 2A2, left). Results were presented as a percentage of the total dendritic length in each quadrant. A similar procedure was used to analyze the wedge from CHPs (CHP wedge analysis, Figure 2A2, right). This analysis was performed using a custom MATLAB (Mathworks) script. In this case, the percentage of the total CHP volume was determined in each quadrant of the anterior and posterior planes.

### 2.3 Shape of CHP

To obtain the actual shape of CHP of an individual neuron, the *convhulln* function in MATLAB was used. The *convhulln* function is based on Quickhull algorithm from Barber et al., 1996 (Barber et al., 1996), which allows the obtain the smallest CHP from a given set of points. The input for the *convhulln* function was the *Segment points-dendrites* data from Neurolucida, which contains the 3D coordinates of each dendritic segment. The outputs of *convhulln* are the list of vertices that form a facet and the volume of the CHP. The volumes of CHPs obtained using *convhulln* coincided with the volumes reported for CH by Neurolucida Explorer (MBF Bioscience).

### 2.4 Intersected volume

Obtaining the volume of the intersected polyhedron required a transformation of the way the CHPs were encoded. For that, using MATLAB, we started from the original CHP and obtained the vertices. By using the *linspace* function, vertices forming each CHP facet were joined using 1000 points per CHP edge. Thus, we ended with a transformed CHP defined by points lined up along edges. To obtain the intersected polyhedron between pairs of CHPs, we proceeded as follows: using the *inhull* function (D’Errico, 2021), the points from each transformed CHP edge that are inside the other CHP of the pair were obtained, and then the *convhulln* function was used to obtain the intersected CHP and the intersected volume.

#### 2.4.1 Intersection ratio

It was reasoned that because the intersected volume depends strongly on the volume of individual neurons CHPs (for instance, the intersected volume of two large but minimally intersected neurons could be the same as the intersected volume of two small but largely intersected neurons), it was necessary to find a ratio that normalized for the volume. As such, it was decided to work with an intersection ratio (IR), that is defined as follows:

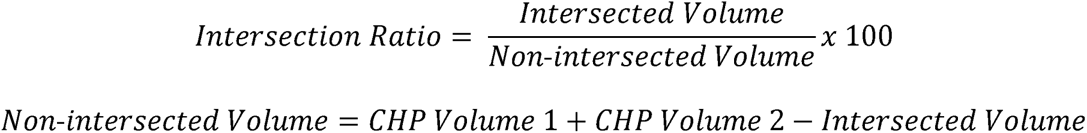

We decided to normalize by the non-intersected volume, and not by the total added volume of both individual neurons, so that the ratio approaches 100 for maximally overlapping CHPs. In any event, the results in the vast majority of analyses did not vary between ratios using either the *non-intersected volume* or the *total added volume* as divisors (not shown), nor when total intersected volume was used.

#### 2.4.2 Dendritic length and nearest neighbor inside intersected volume

First, using the *Segment points-dendrites* data from Neurolucida Explorer in MATLAB, each neuron dendritic segment was resampled, in order that the distance between dendritic segments were equal to 0.001 µm (new dendritic segments). Then, after determining the intersected volume between pairs of CHPs, the *inhull* function was used to search which new dendritic segments from each neuron where inside the intersected volume. Following, the dendritic length inside the intersection (in µm) corresponded to: number of new dendritic segments (inside the intersected CHP) x 0.001.

After determining which dendritic points were inside the intersected volume, the nearest neighbors between the new dendritic segments of the pair of neurons were determined using the *knnsearch* function. Then, the nearest neighbor from the pair of neurons was the minimum value from the output matrix from *knnsearch*.

### 2.5 Methods of CHP geometry modeling

The following methods were developed using custom MATLAB scripts. For each of these methods, IR was calculated at increasing distances between cell bodies (i.e., cell bodies of each neuron, except for normalized cell body, see below), from 0 to 400 µm, at 10 µm steps. One neuron of the pair was moved at these distances across the ML axis, then the DV axis, and finally the AP axis.

#### 2.5.1 Normalized volume

Neurons CHPs were centered in the origin in relation to their cell body, and then the CHPs vertices coordinates were multiplied by the following conversion factor:

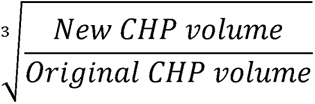

The converted CHPs conserve their shape and the position of their cell body.

#### 2.5.2 Normalized shape

To address the effect of CHP shape on IR, we needed first a rationale and then a method to define an average CHP in 3D. The chosen rationale can be narrated as follows: let’s imagine an apex centered at 0, 0, 0 and from which rays emanate in all directions in space. Then the cell bodies of a pair of CHPs are placed at the apex (as shown in Figure 4C, see also Supplementary Figure 1). Any single ray will intersect the facets of the two CHPs at distances d1 and d2. Then, a point at mid-distance between d1 and d2 is defined, and the exercise is repeated for all rays emanating from the apex. A new polyhedron will be created by all points located at mid-distance between the facets of both CHPs at every solid angle. This procedure can be projected to a larger number of neurons CHP, in which case the points of the new polyhedron will correspond to the average of all *n* distances at which rays intersect the facets of all CHPs in a given ray direction.

Explained in detail, the procedure is based on a surface-to-point transformation procedure. It starts by placing the cell body of a CHP on the center of a Cartesian space (0, 0, 0) where the three orthogonal axes (ML, DV and AP) intersect. Then, a plane (e.g. AP – ML plane) orthogonal to one axis (DV) is chosen, and the distances from the center to the CHP facets in that plane are measured. This is repeated 360 times in 1-degree steps, in an anticlockwise direction. That defines a polygon formed by 360 points (Supplementary Figure 1 A1-5). To move from that polygon to a 3D polyhedron, the original plane is systematically tilted in 1-degree steps around a new axis (e.g. the ML axis). That creates a polyhedron with 64800 points on its surface (Supplementary Figure 1 A6-10). This process for the ML-DV plane, and for the AP-DV plane. To obtain a CHP average from a given number of CHPs, the transformed CHPs were centered in the origin, using the cell body as a reference point. Then, the mid-distance between analogous points on the surface of their respective CHP was determined. A simplified 2D explanation for the average process of 2 or 15 convex hull polygons is provided in Supplementary Figure 1 A and B, respectively. It should be noticed that a consequence of this procedure is that the resulting polyhedron (or resulting polygon in the simplified 2D explanation of Supplementary Figure 1 A and B) is likely not to be convex all around its surface (Supplementary Figure 1 A), for which an approximate convex hull must be defined using the *convhulln* function. On the other hand, the more CHP are included, the more similar is the resulting polyhedron to its convex hull (Supplementary Figure 1 B).

#### 2.5.3 Normalized cell body position

First neurons CHPs were centered in the origin. Then, the centroid of each CHP was obtained using the *centroid* function (Michael Kleder, 2022). Following, the CHPs were centered to the origin subtracting the centroid coordinates to the corresponding vertices coordinates. With these, the cell body of the neurons CHPs were changed to the centroid.

#### 2.5.4 Average neuron according to isotropy index, morphology, and dendritic distribution

The isotropy index was used to estimate how different from a sphere each CHP was. Each neuron CHP was centered in the origin (coordinate 0,0,0), using the cell body as the reference point. Then, a sphere, that had the same volume as the CHP, was also centered in the origin (in relation to the sphere centroid). Following, the intersection volume was obtained. The isotropy index correspond to the ratio between the intersected volume and the CHP volume. To obtain the average isotropy index neuron, the median of isotropy index from the population was calculated.

For the average morphology neuron, the following parameters obtained from Neurolucida Explorer were used: dendritic length, dendritic tree number, maximum dendritic order, number of segments and convex hull volume. For each parameter, the absolute deviation from the mean was obtained (absolute value of the subtraction between each data and the mean of the corresponding parameter). Then, for each neuron, the sum of the absolute deviations from the mean was calculated. The neuron with lower value corresponded to the average morphology neuron.

For average wedge neuron, the dendritic wedge data for the 4 anterior and 4 posterior quadrants of each neuron was obtained. Then, the absolute deviation from the mean was calculated for each quadrant. Following, for each neuron, the sum of the absolute deviations from the mean was calculated. The neuron with lower value corresponded to the average wedge neuron.

### 2.6 Computational and statistical analysis

MATLAB R2018a was used on a Lenovo P51 (Intel i7-7820HQ, 16 Gb of RAM, NVIDIA Quadro M1200) with Windows 10. Statistical analyses were performed on MATLAB. Two-way ANOVA was used (Figure 2A3). Spearman correlation was used for Figure 2A1, B3. Friedman test followed by Dunn-Sidak post-test was used (Figure 8C). Wilcoxon rank sum test and signed rank test, with Sidak multiple comparisons correction (*pval_adjust* function, (Fachada & C. Rosa, 2018)) were performed (Figures 6, 7, 8). An exponential decay model was used (Figure 8B). Significance for all statistical tests was set at p < 0.05.

## 3. RESULTS

### 3.1 CHP volume and intersection between CHPs pairs: method and validation

In this work, we developed methods to determine the intersection between the dendritic domain of pairs of adjacent neurons and further studied how the distance between neuronal pairs, as well as the volume, shape and cell body position of each respective dendritic domain affects how much their domains intersect. For most of this work, we used a sample of 15 previously reconstructed VTA DA neurons (Figure 1B, original data from (Montero et al., 2021)). VTA DA neurons show a high overlap of their dendritic fields (Montero et al., 2021; Phillipson, 1979), making them a useful sample to test dendritic field intersection using CHPs.

**Figure 1.**
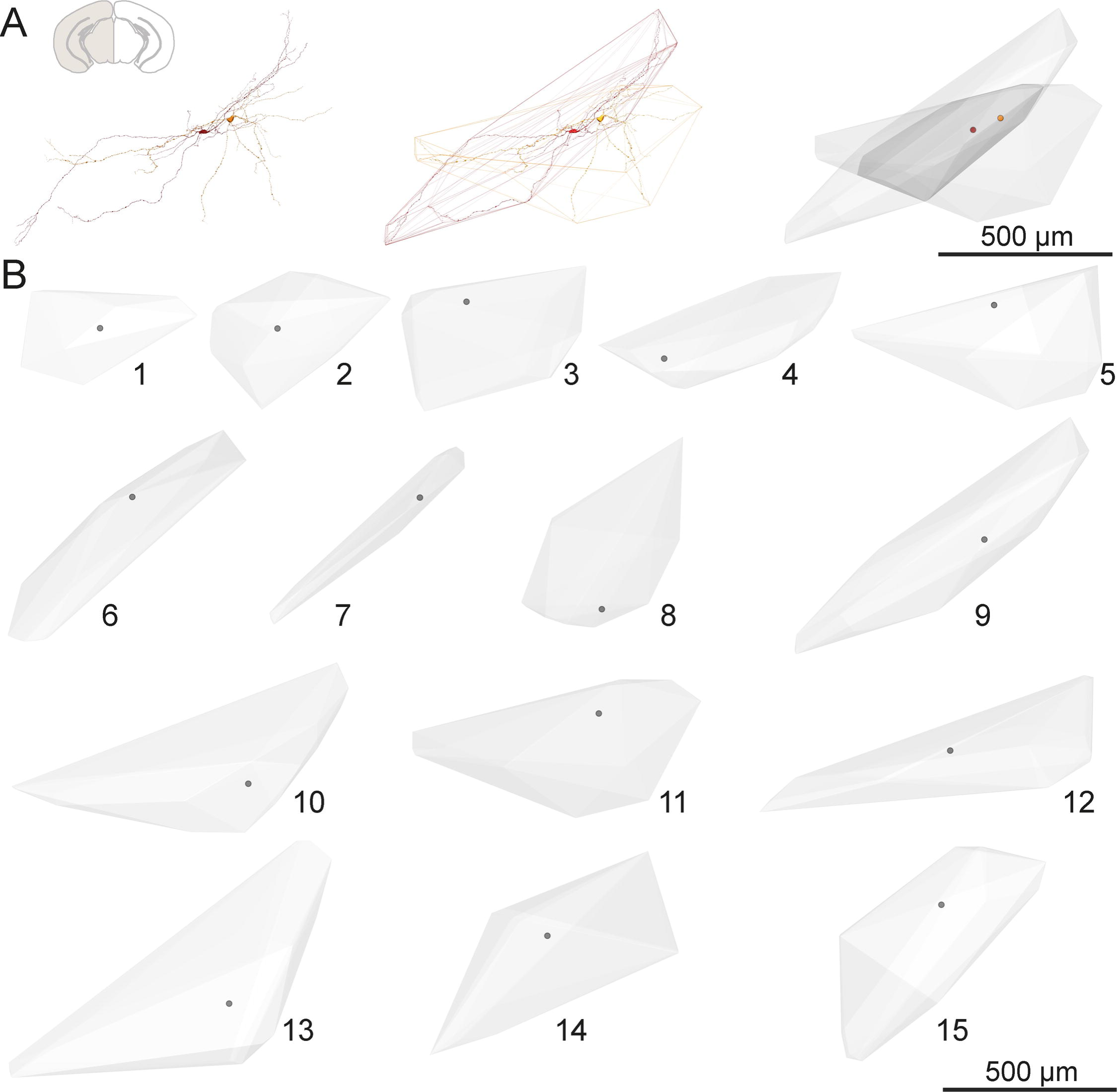
Dendritic domain intersection. A) A pair of reconstructed VTA DA neurons (left) and their corresponding CHPs (center). The intersected volume is shown in darker gray volume (right). Cell bodies are represented with discs. B) All CHPs of VTA DA neurons used in this study (n = 15, frontal view only), numbered. Cell bodies are represented with discs.

**Figure 2.**
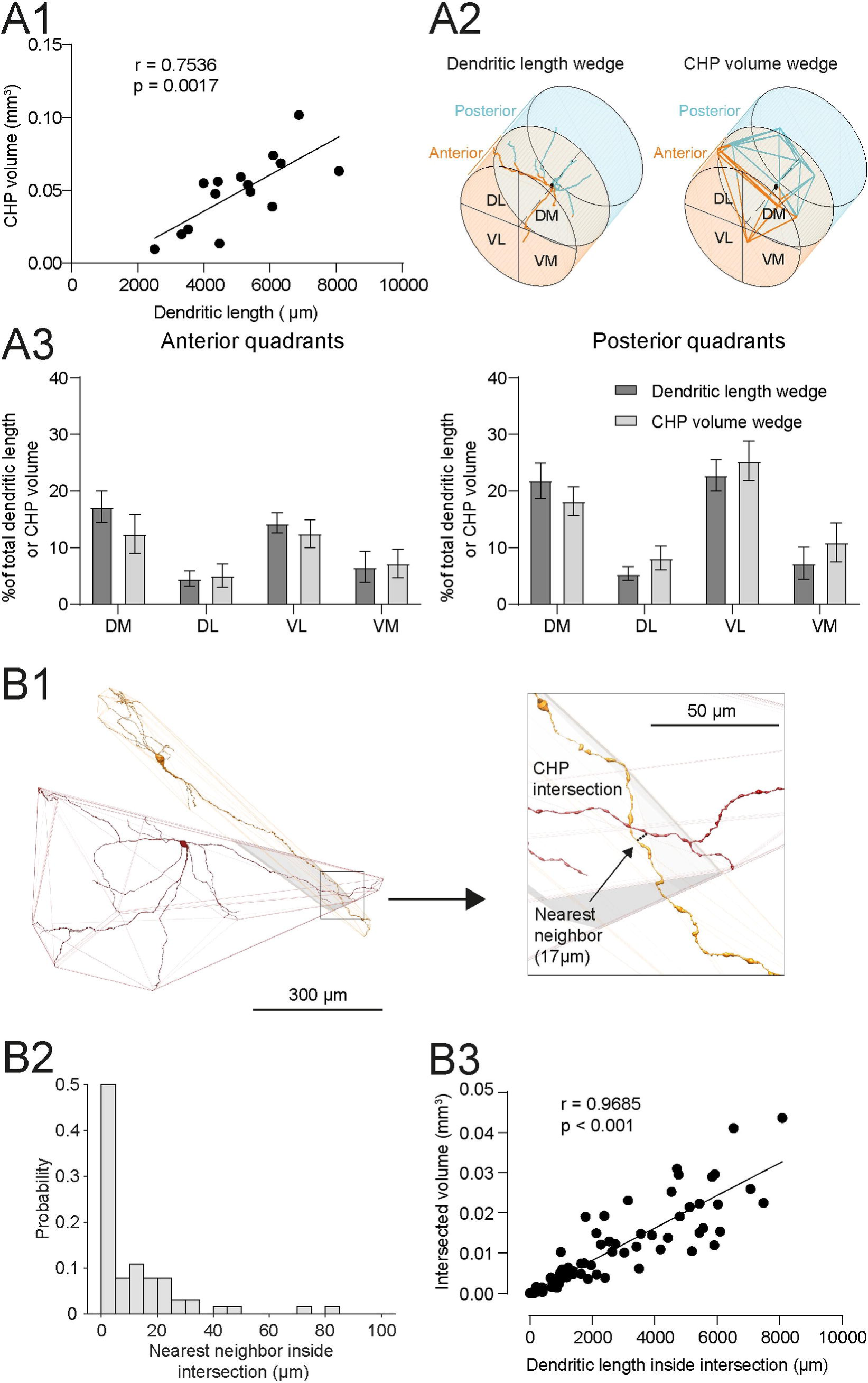
Validation of CHP intersection. A1) Correlation between CHP volume and dendritic length from a sample of VTA DA neurons (n = 15). Spearman correlation was used. Line represents a linear regression. A2) Schematic illustrating the 3D wedge analysis, for dendritic length (left) and CHP volume (right). Dendritic length was measured in each quadrant and normalized using the total dendritic length of the neuron. For the CHP wedge analysis, the volume of the CHP in each quadrant was measured and normalized by the total CHP volume. Dendrites or CHP wedges inside the anterior or posterior plane were colored orange or cyan, respectively. A3) Distribution of dendritic trees and CHP volume for VTA DA neurons (n = 15 neurons), in the anterior (left) and posterior (right) quadrants. When a two-way ANOVA was performed, no significant differences were found: Interaction: Anterior quadrants, F_(3,42)_= 0.8531, p=0.4729; Posterior quadrants, F_(3,42)_= 0.8369, p=0.4813. Bars represent mean ± SEM. B1) Representative example of a pair of VTA DA neurons whose CHPs intersect (left, mirrored frontal view, reflected for layout porpuses). Inside the intersected volume, dendrites from both neurons are found (right). The nearest neighbor distance between dendritic points of the pair of neurons is depicted (black dotted line). B2) Distribution of nearest neighbor values between pair of dendrites inside CHP intersection (n = 64 pairs of CHPs). B3) Correlation between the intersected volume and the dendritic length inside intersection (n = 64 pairs of CHPs). Spearman correlation was used. Line represents a linear regression.

A depiction of volume intersection between the dendritic domain of two neurons is shown in Figure 1. The two reconstructed VTA DA neurons were placed in their approximate position within the brain tegmentum (A, left, as shown in (Montero et al., 2021)). When considering the CHP derived from the disposition of dendrites in space (Figure 1, center), an intersected volume was defined (Figure 1, right, gray volume, more details in methods).

We started by validating the use of CHP as a proxy of the distribution of dendrites in space (Figure 2). First, the CHP volume showed a significant correlation with the dendritic length in our sample (Figure 2A1, r = 0.7536, p = 0.0017, Spearman correlation). To better understand this correlation, we checked whether the extension of dendrites (measured in length) in a space portion surrounding the cell body (Figure 2A2, left) was proportional to the CHP extension (measured in volume) in the same space portion (Figure 2A2, right). For that, we performed a wedge analysis (see Methods), dividing the dendritic field of each neuron into 4 anterior and 4 posterior quadrants, as centered on the cell body, and determined the proportion of dendritic length in each of these quadrants. We compared that to the proportion of CHP volume in each respective quadrant (Figure 2A3) and found that there were no significant differences between them (Two-way ANOVA Interaction: Anterior quadrants, F_(3,42)_=0.4797 p=0.6981; Posterior quadrants, F_(3,42)_=0.6734 p=0.5732), thus validating the use of CHP as an adequate proxy for the distribution of dendrites in space.

We then evaluated the proximity between dendrites of each neuron in the CHP intersection. Figure 2B1 shows a representative pair of VTA DA neurons, located in their respective positions in the VTA (Montero et al., 2021), with their corresponding neuronal CHPs and the CHP intersection (gray). It can be observed that both neurons showed dendrites inside the CHP intersection. Relevantly, the distance between the nearest neighbor segments was 17 µm. When this was assessed for all the VTA DA neuronal pairs in their respective position in the VTA (more details about this in Figure 8 and in methods), a left-skewed distribution was observed, with a median value of 4.6 µm (Figure 2B2, n = 64 pairs with an IR>0), indicating that inside the CHP intersection dendrites are close to each other.

We found that for 84% of the pairs, nearest neighbor distances lower than 25 µm were observed. Moreover, a significant correlation between the intersection volume and the dendritic length inside the intersection was found (Figure 2B3, r = 0.9685, p < 0.001, Spearman correlation). Overall, these results show that the intersection of neurons CHPs is a representative proxy of the intersection between dendritic fields.

### 3.2 CHP intersection dynamics: modeling the influence of CHP volume, shape and cell body position

Following method validation, we evaluated how the intersection between pairs of neurons CHPs changes when the distance between CHPs increases (Figure 3A). We used an intersected ratio (IR, more details in methods) as a normalized index of intersection between pairs of neurons CHPs. Then, IR values were obtained from pairs of VTA DA neurons, first with their cell bodies aligned (distance between cell bodies = 0 µm) and then after one of the neuron CHP was moved across the ML, DV or AP axes (in 10 µm steps, until 400 µm). In Figure 3B we show some representative examples of IR vs distance curves. We found a high variability in IR as a function of distance. For instance, in Figure 3B (ML), the green curve shows a pair of CHPs whose IRs first increased, then reached a peak, and then decreased, in contrast to the magenta and red curves, whose IR values only decreased. These results indicate that IR between neuronal pairs depends on the distance between their cell bodies, but also on other geometrical factors.

**Figure 3.**
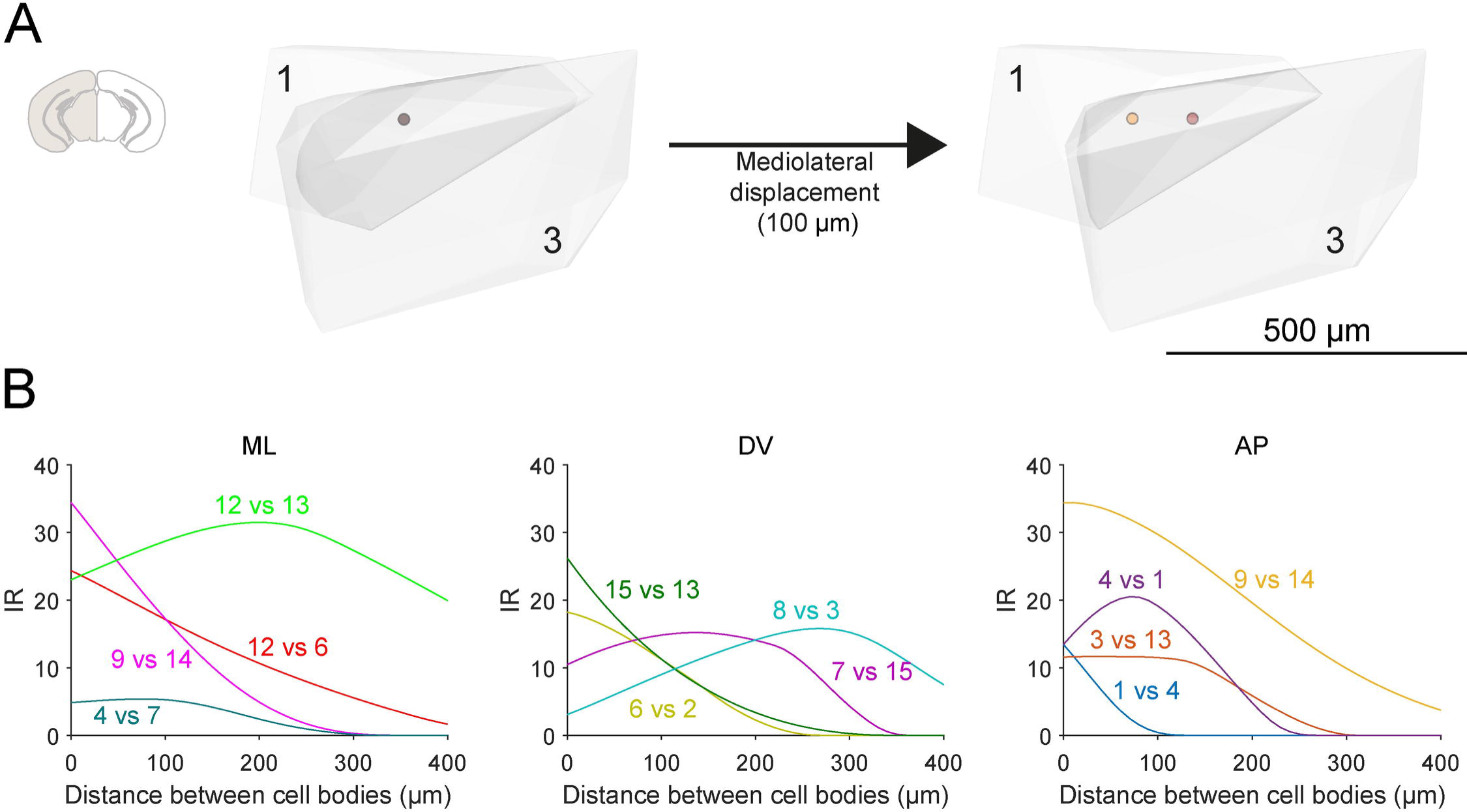
CHP intersection dynamics. A) Example of a pair of VTA DA neurons CHPs (frontal view, numbers relate to Figure 1B) that were initially centered in their cell bodies (black disc) and following the displacement of one of the neuron CHP (3) across the ML axis. Intersected volume is shown in darker gray. Cell bodies are represented with discs. B) Representative examples of IR analysis between all pairs of VTA DA neurons across the ML, DV and AP axes. IR was tested from 0 to 400 µm, in 10 µm steps. In this Figure, depicted pairs are indicated by their respective numbers in Figure 1B).: First number corresponded to the neuron CHP that stayed in a fixed position, second number corresponded to the neuron CHP that was moved across the axis. Note the diversity of intersection dynamics observed.

**Figure 4.**
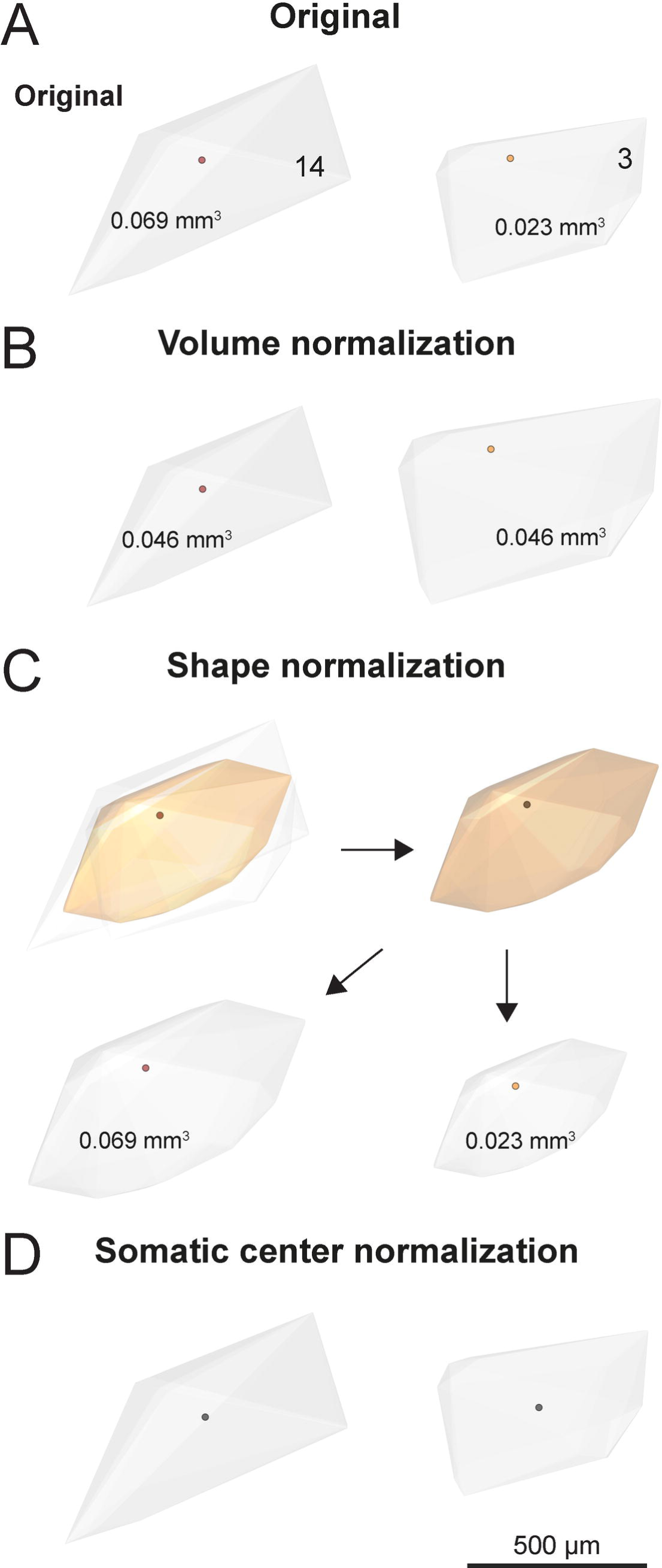
Modeling of neuron CHPs intersection dynamics. A) Original, unmodified pair of VTA DA neurons CHPs. Numbers inside are related to Figure 1B. CHP volume is depicted inside each CHP. Cell bodies are represented as discs. B) Normalized volume modeling: in a given population of neurons CHPs (A), the average CHP volume was obtained (in this case, the average of the pair, 0.046 mm^3^) and then all the neurons CHPs were normalized to that average CHP volume. C) Normalized shape modeling: first, neurons CHPs were centered, using their cell bodies as reference (represented as discs). Then, the normalized shape was obtained (more details in methods and Supplementary Figure 1). Following, new CHPs were obtained from this normalized shape, using the original CHPs volumes of the neurons. D) Normalized cell body position: in a given population of neurons CHPs (A), the centroid of each CHP was obtained and used as the reference for that CHP (black discs).

To understand how the differences in geometry between neuronal pairs affect intersection, we separately modeled the influence of neurons CHP volume, shape, and cell body position (Figure 4, see methods) by running the same previous simulations (as in Figure 3), but this time with all pairs having either a normalized volume, a normalized shape or a normalized cell body position. More precisely, we first transformed all CHPs to a normalized volume, which was equal to the population mean for VTA DA neurons (0.047 mm^3^). In Figure 4A,B we show a simpler description of the procedure using only two neurons CHPs. Their respective volumes are 0.069 mm^3^ and 0.023 mm^3^ (Figure 4A), therefore one of them decreases its volume to the average value (Figure 4B, neuron CHP 14), and the other increases its volume to the average value (Figure 4B, neuron CHP 3). Note that the CHP shape remains unchanged with this procedure.

To examine the effect of differences in CHP shape on IR we reasoned that, similarly, this could be assessed using a normalized shape from the population of CHPs. We set to find a CHP that we defined as the CHP of the polyhedron formed by the points located at average distances from the surfaces of all CHP, with their cell bodies aligned. After the normalized CHP is defined, its volume can be modified to the volume values of the original neurons CHPs. In Figure 4C we show a simpler description of the procedure using only two neurons CHPs. The CHPs of the previous pair now were combined to generate a normalized shape following the averaging rules already explained (Figure 4C, top row, more details in Supplementary Figure 1 and methods). Then, the new normalized CHP was given back the original CHP volumes (Figure 4C, bottom row).

The normalized shape of all VTA DA neurons is shown in Figure 5A. As it can be appreciated, the average shape exhibits the essential geometrical features of this dopaminergic population, particularly the dendritic domain size and the extension in the ML, DV and AP axes. In frontal view, we observe that the CHP was oriented dorsomedially to ventrolaterally, with the cell body at mid position, in agreement with the distribution of VTA neurons dendritic field, as described in (Montero et al., 2021).

**Figure 5.**
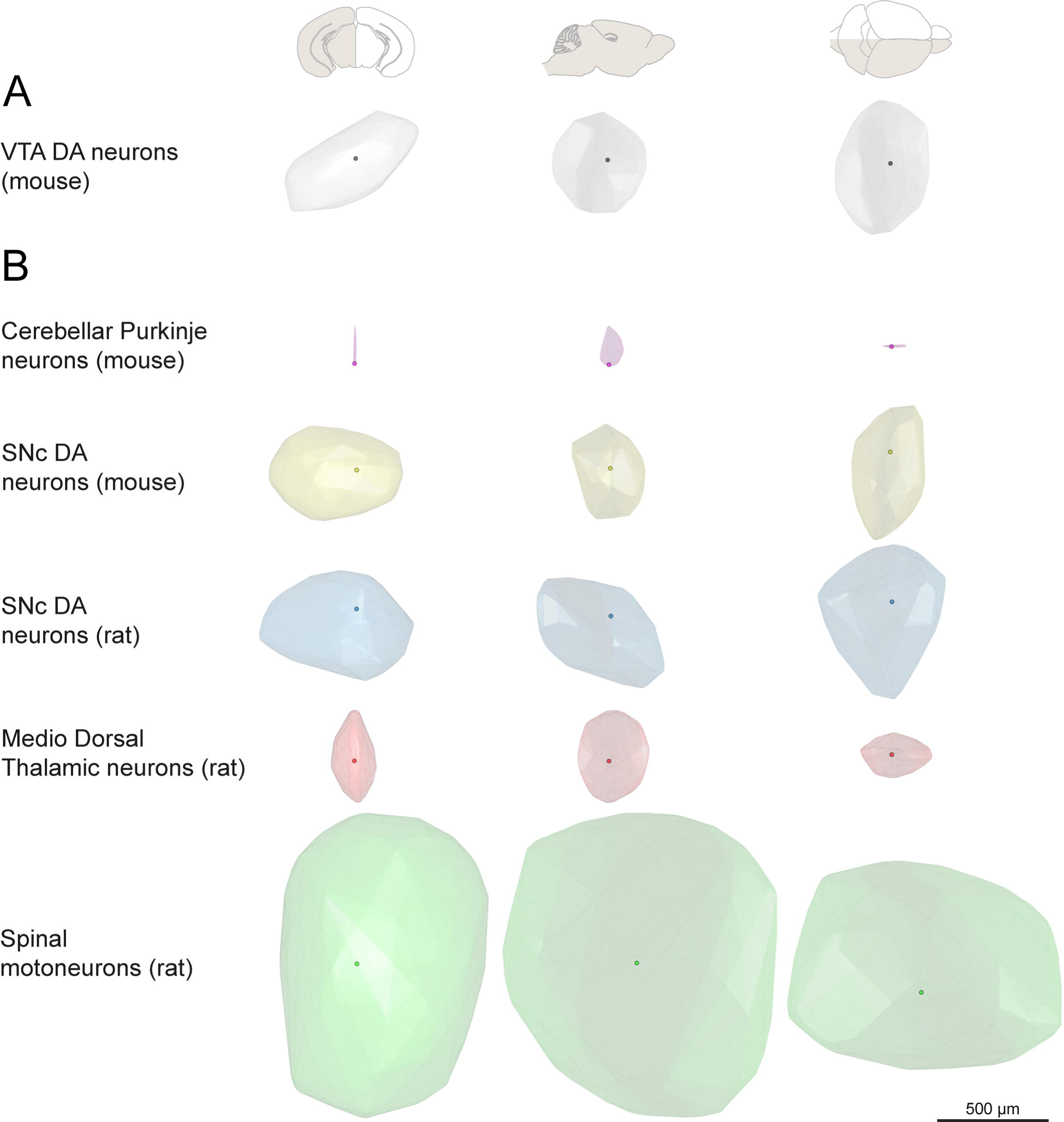
Normalized CHP shape for different neuronal types. The normalized CHP shape was obtained for: A) VTA DA neurons (mouse); B) cerebellar Purkinje neurons (PC, mouse), SNc DA neurons (mouse and rat), mediodorsal thalamic neurons (MD, rat) and spinal motoneurons (MN, rat). The volume for each normalized shape was modifed to match the average CHP volume of the corresponding sample. Neurons are shown from the frontal, lateral and dorsal views. Cell bodies are represented with discs.

**Figure 6.**
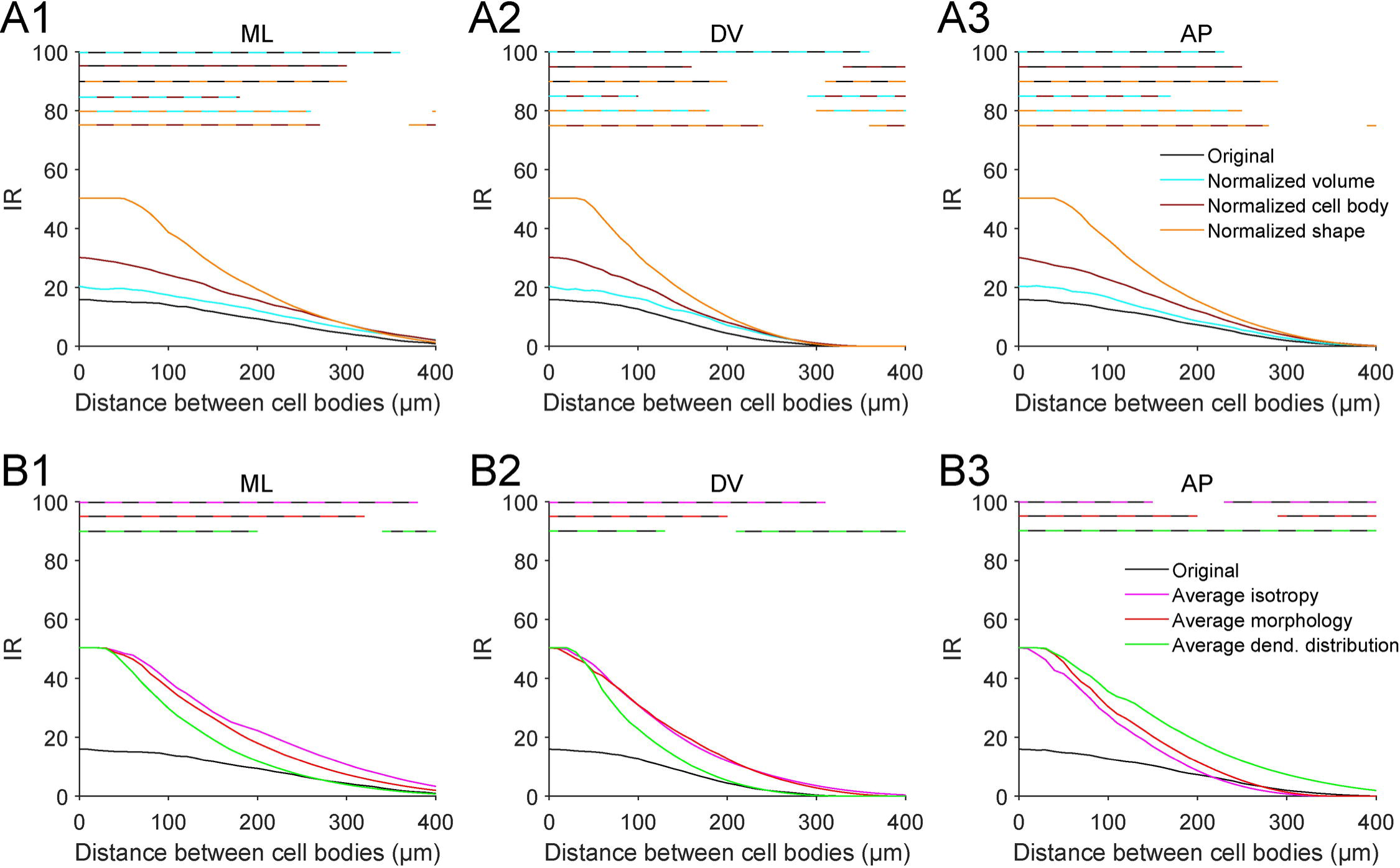
CHP modeling for VTA DA neurons. A1-3) IR vs distance plots across the ML (A1), DV (A2) and AP (A3) axes were performed for all the possible pairs of VTA DA neurons (210 pairs per distance) and compared between modeling conditions. IR was measured between 0 to 400 µm (10 µm steps). Discontinuous color horizontal lines on the top of each plot depict the significant differences (p < 0.05, Wilcoxon signed rank test with Sidak multiple comparisons correction) between original vs normalized volume (black and cyan line), original vs normalized cell body (black and red line), original vs normalized shape (black and orange line), normalized volume vs normalized cell body (cyan and red line), normalized volume vs normalized shape (cyan and orange line), and normalized cell body vs normalized shape (red and orange line). Solid line represents the median. Error bars are omitted (data with error bars are shown in Supplementary Figure 2). B1-3) Comparison between normalized shape modeling for VTA DA neurons vs average neurons according to isotropy index, morphology, and wedge dendritic distribution analysis (details in methods). Discontinuous color horizontal lines on the top of each plot depict the significant differences (p < 0.05, Wilcoxon signed rank test with Sidak multiple comparisons correction) between original vs average isotropy index (black and magenta line), original vs average morphology (black and red line), and original vs average wedge (black and orange) neurons. Solid line represents the median. Error bars are omitted.

**Figure 7.**
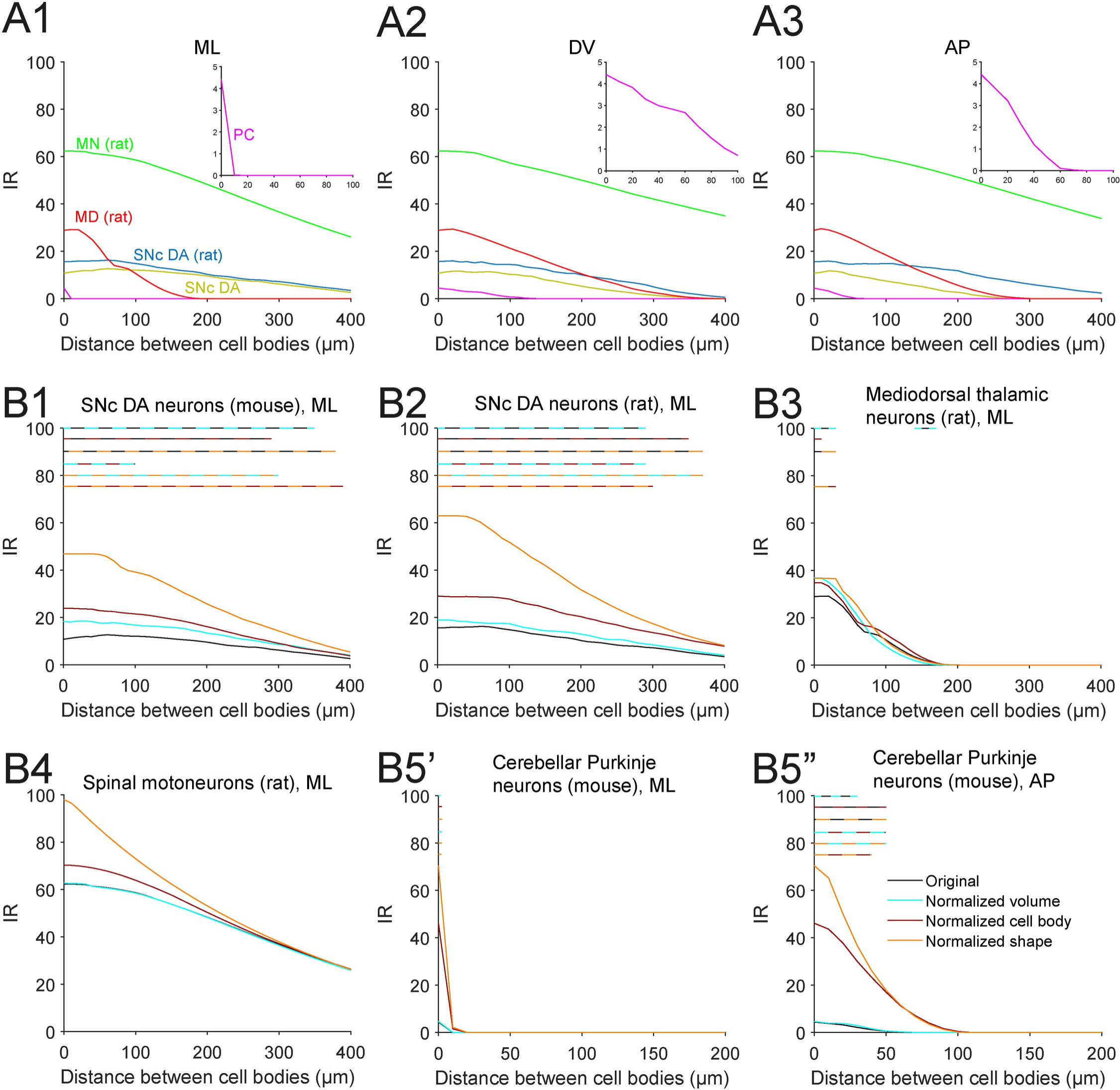
CHP modeling: examples from different neuronal types. A1-3) Original data analysis: IR vs distance plots across the ML (A1), DV (A2) and AP (A3) axes were performed for all the possible pairs of SNc DA neurons (mouse and rat), cerebellar Purkinje neurons (PC, mouse), mediodorsal thalamic neurons (MD, rat) and spinal motoneurons (MN, rat). Solid lines represent the median. Error bars were omitted. B1-6) Within region modeling analysis: IR vs distance plots are shown for modeling conditions of: SNc DA neurons (mouse), ML axis (B1); SNc DA neurons (rat), ML axis (B2); Mediodorsal thalamic neurons (rat), ML axis (B3), spinal motoneurons (rat), ML axis (B4), cerebellar Purkinje neurons (mouse), ML (B5’) and AP (B5’’) axes. IR was measured between 0 to 400 µm (10 µm steps). Discontinuous color horizontal lines on the top of each plot depict the significant differences (p < 0.05, Wilcoxon signed rank test with Sidak multiple comparisons correction) between original vs normalized volume (black and cyan line), original vs normalized cell body position (black and red line), original vs normalized shape (black and orange), normalized volume vs normalized cell body position (cyan and red line), normalized volume vs normalized shape (cyan and orange line), and normalized cell body position vs normalized shape (red and orange line). Solid line represents the median. Error bars are omitted.

**Figure 8.**
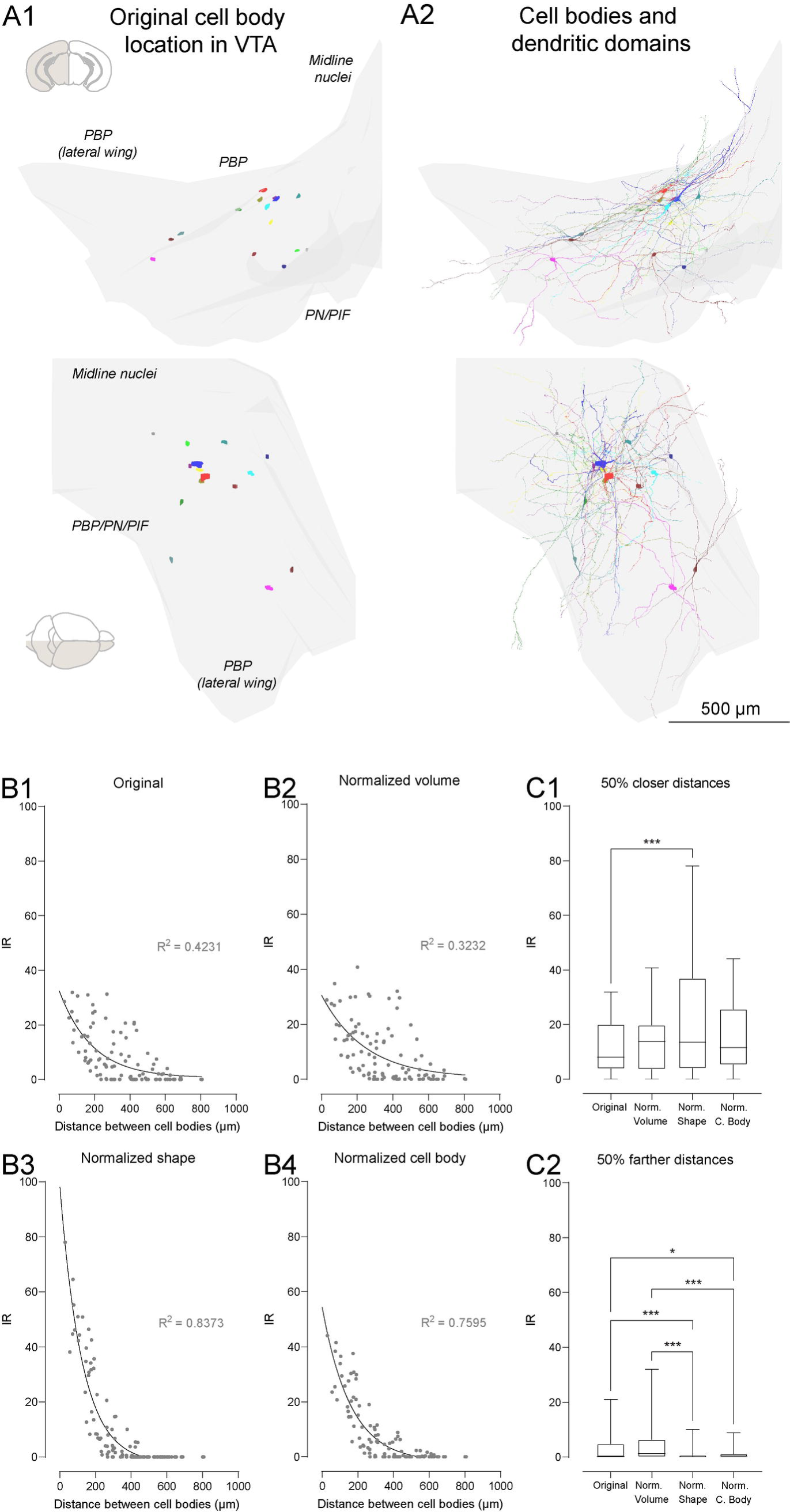
CHP intersection analysis: VTA DA neurons in their original brain position. A1-2) Rendered 3D map of the VTA, showing the cell bodies (A1) and cell bodies plus their dendrites (A2) of the 15 VTA DA neurons, as in their real positions in the VTA (determined in (Montero et al., 2021)), and seen in frontal (first row) and dorsal (second row) views. B1-4) IR vs distance plots of the 105 possible pairs of VTA DA neurons, as tested unmodified (original, B1) and with CHP normalized volume (B2), normalized shape (B3), and normalized cell body location (B4). Each point represents the IR value of a pair of CHPs. Black line corresponds to an exponential decay non-linear regression. C1-2) IR values from (B) were grouped according to their distance between cell bodies: 50% closer distances (C1, n = 53) and 50% of farther distances (C2, n = 52) between cell bodies. Data are represented as box and whiskers, showing the median (middle line) and the 25th and 75th percentile in the box and the lowest and highest values (whiskers). * p < 0.05, *** p < 0.001 using a Friedman test followed by a Dunn’s multiple comparisons test.

Finally, we took into account the fact that, as it can be noted in Figure 1B, the cell bodies of real VTA DA neurons are normally displaced from the center of the dendritic domain. We considered this a relevant factor for intersection and therefore defined a third normalization procedure, in which the cell body of each CHP was moved to the CHP’ centroid. Again, in Figure 4D we show a simpler description of the procedure using only two neurons CHPs (more details in methods).

We applied the modeling procedures described above to our population of VTA DA neurons (Figure 6). In each condition and in the ML, DV or AP axes, all the possible pairs of VTA DA neurons were analyzed (210 pairs in total, using repeating pairs given that a neuron “a” in a pair is first tested moving away from neuron “b”, and then neuron “b” is tested moving away from neuron “a”). Compared to the original data (no changes in CHP geometry), a significant increase in IR was found in the normalized volume condition across the ML (Figure 6A1), DV (Figure 6A2) and AP (Figure 6A3) axes (p < 0.05 using Wilcoxon signed rank test with Sidak multiple comparisons correction, more details of statistical differences between conditions in Figure 6A), but the magnitude of change in IR was low. For the normalized cell body position condition, significantly higher IR values were observed in the three axes, compared to the original data and also to the normalized volume condition (Figure 6A1-3). However, the modeling condition that showed the largest effect on IR values across each axis, was the normalized shape condition (Figure 6A1-3). These results indicate that differences in IR values between pairs of VTA DA neurons CHPs are influenced, in decreasing order, by the CHP shape, cell body position, and volume.

To give further sustain to the results describing the influence of shape normalization on intersection (see discussion), we used CHPs from neurons that exhibited average values for morphology, isotropy and dendritic distribution characteristics, as follow: 1. average isotropy neuron: for each neuron CHP, an isotropy index was calculated to determine how much a CHP is similar to a sphere of the same volume (more details in methods); then the median population value was found (0.46), and the neuron with the closest value was selected (in this case neuron 7 (Figure 1B)); 2. average morphology neuron: using the dendritic length, dendritic tree number, maximum dendritic order, number of segments and convex hull volume, we found the neuron with the lower absolute deviation of the mean from the sample (more details in methods), which corresponded to neuron 14 (Figure 1B); 3. average dendritic distribution neuron: using the dendritic wedge data for the 4 anterior and 4 posterior quadrants (Figure 2A3), we found the neuron with the lower absolute deviation from the mean from the sample (more details in methods), which corresponded to neuron 1 (Figure 1B). Each average neuron was then analyzed using the same approach as for the normalized shape analysis: starting with one of the three average neurons, we created 15 new CHPs, all with the average neuron CHP shape, and then gave to each new CHP the respective 15 volumes from the VTA DA neurons population. And then determining IR values between 0 to 400 µm (10 µm steps) across the ML, DV and AP axes.

Figure 6B shows the IR curves for average isotropy neuron, average morphology neuron and average dendritic domain neuron. Each condition was compared with the original data analysis across each axis. All three average neurons showed significantly higher IR values than the original condition (p < 0.05 using Wilcoxon signed rank test with Sidak multiple comparisons correction, more details of statistical differences between groups in Figure 6B) and almost identical magnitudes to that of CHP shape normalization in 6A.

We wanted to further examine the diversity of distance versus IR curves shown previously (Figure 3), this time focusing on the types of decay curves observed. For the original data, we found that 51% of the curves for all pairs and axes tested, described an initial decay (“decay curves”, as the magenta or red curves in Figure 3B, ML), whereas 49% of curves exhibited an initial rise, a peak, and then a decay (“peak curves”, as the green curve in Figure 3B, ML). After volume normalization, the proportion of decay and peak curves remained similar to the original data (51% vs 49%). Interestingly, in the normalized cell body condition, an increase in decay curves cases was observed (57%, versus 43% of peak curves). Yet the larger effect was found in the shape normalization condition, where peak curves disappeared and thus 100% of the curves ended as decay curves. Overall, these results are in line with the previous findings of this section, highlighting that CHP shape is the main factor that contributes to the variability in intersection dynamics.

### 3.3 CHP intersection dynamics: examples from other neuronal types

To further characterize our proposed method to study intersection of dendritic fields, we searched for in vivo labeled neurons in the neuromorpho.org database (more details in methods). The following pools of neurons were used: cerebellar Purkinje neurons (PC, n= 10, (Nedelescu et al., 2018)), substantia nigra pars compacta (SNc) DA neurons from rat (n = 10, (Henny et al., 2012)) and mouse (n = 15, (Farassat et al., 2019; Meza et al., 2018; Montero et al., 2021)), medio dorsal thalamic neurons (MD, n = 5, (Kuramoto et al., 2017)) and spinal motoneurons (MN, n = 3, (Rotterman et al., 2014)). Figure 5B shows the normalized shape of these neuronal populations.

We first computed the IR dynamics of the neuronal CHPs (original data) of each region, and qualitatively examined differences between cell types (Figure 7A). MN neurons show higher IR values in the three axes, followed by MD, SNc rat and mouse neurons, and PC cells, although each population shows specific differences in each axis.

As with VTA neurons (Figure 6), we modeled CHP geometry for PC, SNc, MD and MN neurons populations, using the normalized volume, normalized shape and normalized cell body position conditions (Figure 7B). We show the most representative axis of each neuronal population. As in the original condition analysis (Figure 7A), mouse and rat DA neurons showed similar curve profiles after the application of each modeling conditions in the ML axis (Figure 7B1-2). Specifically, in both SNc mouse (Figure 7B1) and rat (Figure 7B2) analysis, normalized volume showed increased IR values than the original condition (p < 0.05, Wilcoxon signed rank test with Sidak multiple comparisons correction, more details of statistical differences between conditions in Figure 7B1-2), but the magnitude of change in IR is low. The normalized cell body location modeling also showed significantly higher values than the original and also the normalized volume conditions in both SNc mouse and rat neurons, but the magnitude of change was higher in SNc rat. The largest effect for both mouse and rat SNc DA neurons was observed in the normalized shape condition, which was significantly higher than the other three conditions. These results were very similar to those found with VTA DA neurons (Figure 6A), indicating a common underlying dendritic organization.

Interestingly, modeling conditions applied on MD neurons only significantly modified IR values on short distances on the ML axis (p < 0.05, Wilcoxon signed rank test with Sidak multiple comparisons correction, more details of statistical differences between conditions in Figure 7B3). These results indicate that neither volume, shape or position of the cell body is relevant to explain the intersection dynamics in MD neurons.

For MN, no significant differences were observed between conditions, although normalized shape showed a trend of higher values than the other conditions (Figure 7B4). The lack of significant differences could be related to the low sample of neurons in this region (3 neurons, 6 pairs analyzed at each distance).

Finally, for PC, we first show the ML axis as a positive control for our results, considering the mainly 2D structure of the PC dendritic field, and minimal extension in the ML axis. In this sense, the modeling conditions only showed an effect at a distance between cell bodies of 0 µm, with a fast decrease of IR values at 10 µm (Figure 7B5’). A different result was found at the AP axis (Figure 7B5’’), a more biologically relevant condition, considering the default sagittal arrangement of these neurons in the cerebellum. A significant difference between the original and normalized volume condition was observed, but the magnitude of the difference was low (p < 0.05, Wilcoxon signed rank test with Sidak multiple comparisons correction, more details of statistical differences between conditions in Figure 7B5’’). Higher IR values were found after normalizing cell body location and shape, until 50 µm. In summary, these results indicate that intersection dynamics appear to be cell-type specific.

### 3.4 CHP intersection dynamics using neurons real position in the VTA

As a final approach to characterize the uses of the CHP intersection method for neurons, we again used our sample of VTA DA neurons, but now to analyze IR dynamics using the real positions of the neurons in the region (as determined in (Montero et al., 2021)). As seen in Figure 8A, our sample of DA neurons is located in the central region of the VTA (see Discussion). IR values for all the possible pairs were obtained (105, without repeating). This analysis was performed for the original data (Figure 8B1), and the three modeling conditions used in the previous sections: normalized volume (Figure 8B2), normalized shape (Figure 8B3), and normalized cell body location (Figure 8B4). For the original data, high variability in IR values across distance was found (Figure 8B1), as data show a low R^2^ for an exponential decay non-linear regression (R^2^ = 0.4231). A similar IR profile was found in the normalized volume condition (Figure 8B2), confirming a low effect of volume variability in intersection dynamics (as also found in Figure 6A), with an even lowest R^2^ for an exponential decay fit (R^2^ = 0.3232). Interestingly, shape and cell body location normalization drastically changed the data curve profile and reduced the variability of IR values across distances (Figure 8B3-4), as evidenced by a higher exponential decay fit (normalized shape: R^2^ = 0.8373; normalized cell body: R^2^ = 0.7595). Finally, we analyzed the IR values of the closer 50% (0 to 340 µm, n = 53 pairs, Figure 8C1) and farther 50% (340 to 807 µm, n = 52 pairs, Figure 8C2) of distances between cell bodies. For the closer half of distances (Figure 8C1), we found significantly higher IR values in the normalized shape conditions compared to the original data (p < 0.001, Friedman test followed by a Dunn’s multiple comparisons test). On the other hand, for larger distances (Figure 8C2), original and normalized volume conditions were significantly higher than both normalized shape and cell body conditions (details in Figure 8C2).

Overall, the results using real neurons positions in the VTA confirm the principles described in Figure 6: CHP shape, and to some extent cell body location, in VTA DA neurons are the most relevant factors underlying CHP intersection between pairs of VTA DA neurons.

## 4. DISCUSSION

### 4.1 Use of CHPs to study dendritic domain intersection

CHPs have been used to characterize and classify neuronal populations shape or size (Felix et al., 2013; Gärtner et al., 2004; Gertler et al., 2008; Ito, 2020; Kawa et al., 1998; Malmierca et al., 1995; Mihaljević et al., 2015; Rojo et al., 2016). We adapted previously developed computational tools to determine intersected volume between pairs of polyhedrons (Chazelle & Dobkin, 1987; Govender et al., 2018; Muller & Preparata, 1978) to explore in greater detail the effects of various 3D geometrical factors on CHPs intersection, at the single cell level.

The use of neurons’ CHPs to represent dendritic domain size or shape (Felix et al., 2013; Gärtner et al., 2004; Gertler et al., 2008; Ito, 2020; Kawa et al., 1998; Malmierca et al., 1995; Mihaljević et al., 2015; Rojo et al., 2016) poses the question of the extent to which the CHP is indeed an appropriate proxy, given that the vast majority of space within a neuronal CHP is not occupied by dendritic segments. And, similarly, whether intersection between CHPs reflects dendritic domain intersection in any biologically meaningful way. We examined the shape assumption by testing whether CHP and dendritic length space distribution were comparable. We first confirmed that larger CHP does indeed contain more dendritic length (Figure 2A1). Then we examined the spatial distribution of CHP volume and dendritic length and found that they were indistinguishable from each other, as tested using wedge analysis in Figure 2A3. These results support the contention that CHPs are adequate proxies for dendritic distribution. We then tested the CHP intersection assumption by studying the proximity between the dendrites from two neurons located inside the intersected volume. We found that the median nearest neighbor distance between dendrites from different neurons inside the intersected volume was 4.6 µm, and that in almost 85% of the cases, at least one pair of dendrites located at less than 25 µm (Figure 2B2). To put these results in perspective, a 25 µm distance is similar to the longest axis of DA neurons cell bodies (see Figure 1 in (Meza et al., 2018; Montero et al., 2021)). Finally, we determined that the total length of shared dendrites inside the intersected volume tightly correlates with the CHPs intersected volume (Figure 2B3). These data support the assumption that CHP intersection is an appropriate proxy for the intersection of neuronal dendritic fields.

Having confirmed these assumptions, it is important to stress, however, that CHP does not represent and are therefore not useful to describe other aspects of dendritic organization, such as dendritic density or architectural complexity inside the CHP, as CHPs only considers the outer limits of a dendritic domain. Thus, other systematic modeling approaches must be used if one wants to study the influence of dendritic density or architecture on intersection. We consider those possible approaches, which we imagine would require combining methods that model dendritic architecture (Ascoli & Krichmar, 2000; Cuntz et al., 2010, 2012; Takeo et al., 2021) with those developed here to model distance and compute the intersection, of utmost importance and would certainly complement and illuminate the results described in the present study.

### 4.2 Normalized and representative CHPs

In order to study the influence of shape on intersection, an additional challenge in this study was to find a CHP whose shape best represented an otherwise diverse population. We took the direct, though methodologically more challenging approach, to compute the average 3D CHP shape from the population of 15 neurons CHPs. This approach allowed us to model intersection using a normalized shape, while maintaining the original volume and cell body position of CHPs. Now, because the resulting average CHP is smooth and ovoid-shaped, one reasonable concern is that the observed increases in intersection are merely due to these characteristics, and not due to the use of a normalized shape To address this issue, we chose three CHPs whose respective neurons exhibited average morphology (Montero et al., 2021), isotropy (see methods), and spatial dendritic distribution (Figure 2A3 and methods). In this way, we could use CHPs with average characteristics, but whose CHP maintained its pointed, original shape, thus discarding an artificial influence from the smother, ovoid-shaped average CHP.

### 4.3 Role of distance, volume, cell body location and shape on intersection

Our results show that in addition to cell-to-cell distance predictably decreasing intersection, distance versus intersection relationship was not straightforward and had to depend on other geometrical factors. Indeed, intersection was affected by the relative position of each neuron to its pair (i.e. whether one locates besides, above/below, or in front/behind from the other). This is an expected consequence of the dendritic domain of neurons not being spherical (i.e. in that maximal extensions in ML, DV or AP axes differ from each other). The location of the cell body, which we used to identify each neuron and showed displacement from the dendritic domain’s centroid, was also a relevant factor influencing intersection dynamics. Yet perhaps the most noteworthy finding of this study, however, is that dendritic domain intersection is significantly increased by shape homogeneity between neurons (Figure 6-8). Or, said conversely, that dendritic domain exclusion between adjacent neurons is significantly increased by heterogeneity in dendritic domain size and shape.

We found that individual pairs distance vs intersection decay dynamics were diverse. Among them, some curves exhibited a peak, i.e., an initial rise followed by the expected decay. We found this result interesting because it challenges the intuitive assumption that neurons whose cell bodies locate next to each other should have a maximal chance of sharing afferents inputs. We initially assumed that this specific dynamic (peak curves) mostly resulted from the displaced location of the cell bodies within their respective CHP, resulting in dendritic domains sharing little volume with their cell bodies aligned (for instance, in a hypothetical scenario in which two Purkinje-like cells had their cell bodies aligned, but their respective dendritic domains pointing in opposite directions). Interestingly, however, although cell body position normalization diminished the proportion of peak curves, shape normalization completely eliminated them, again emphasizing the critical role of CHP shape in determining intersection dynamics.

### 4.4 Determinants of VTA DA neurons morphology and overlap

The principles that underlie intersection ultimately depend on the size and spatial organization of VTA DA dendrites. As detailed in (Montero et al., 2021), the dendritic tree of individual mouse VTA DA neurons conforms to early descriptions (Kline & Felten, 1985; Phillipson, 1979; Ramón-Moliner & Nauta, 1966) characterizing them by long radiating dendrites, low branching patterns, and overall conforming to the “isodendritic” cell type typical of reticular neurons (Jones, 1995; Ramón-Moliner & Nauta, 1966).

The mechanisms involved in specification of VTA neurons dendritic trees have not been studied in detail yet (Brignani & Pasterkamp, 2017). We speculate that dendritic morphogenesis of VTA DA neurons is likely to rely on the catenin signaling pathway, which plays a critical role in VTA DA neurons neurogenesis and migration (Tang et al., 2009) and, as studied in other brain regions, is essential for dendritic growth and branching (Arikkath, 2009). On the other hand, given the highly overlapping pattern of VTA DA neurons dendritic trees, there must also exist mechanisms that allow the co-existence of dendrites from neighboring cells that belong to the same cell class (dopaminergic). Indeed, in the retina the dendritic domain of neighboring amacrine dopaminergic (and other amacrine) cells overlap extensively. The mechanisms that ultimately instruct or permit dendritic overlap are less known, although evidence points to the self/non-self discrimination capacity of developing neurons which on the one hand, avoid aberrant branching towards themselves, but on the other, permits growing towards other same-class neurons. Interestingly, this is a process mediated by protocadherins (Lefebvre, 2017; Lefebvre et al., 2015), a group of cell adhesion molecules that are also expressed in the VTA (Hertel et al., 2008).

### 4.5 VTA DA neurons orientation and VTA region shape

Brain regions may exhibit structural folding, and therefore same class neurons may exhibit dissimilar orientations. A pertinent question to ask is whether our sampled VTA DA neurons may exhibit different orientations, perhaps related to the natural folding of the VTA, which may have affected the intersection dynamics described here. As seen in the wedge analysis of dendritic length spatial distribution in Figure 2, our neuronal sample exhibited a dorsomedial to ventrolateral preferred orientation, something also evident in the overall orientation of CHPs in Figure 1, and also in the depiction of neurons within the VTA in Figure 8. Therefore, we think that modeling intersection in this sample is attainable. Indeed, the preferred orientation of VTA

DA neurons coincides with the orientation of the VTA nuclei in which these neurons locate, which corresponds to the main parabrachial pigmented (PBP) and underlying paranigral and parainterfasciular (PN/PIF) nuclei (Montero et al., 2021) all of which exhibit a dorsomedial to ventrolateral orientation (Brignani & Pasterkamp, 2017; Fu et al., 2012; Montero et al., 2021), and locate between the VTA midline nuclei (which exhibit a vertical orientation), and the lateral wing of the PBP (which exhibit a dorsolateral to ventromedial orientation). Interestingly, the orientation of VTA DA neurons and VTA nuclei described also coincides with the orientation of migrating dopaminergic neurons precursors exhibit during the final stages of the ventral midbrain development, indicating that the adult dendritic tree retained the orientation that neurites present during development (Bodea et al., 2014; Brignani & Pasterkamp, 2017; Kawano et al., 1995).

Finally, the similar orientation between VTA DA neurons dendritic domain and the VTA may also explain why our modeling approach applied to neurons in their real brain coordinates (Figure 8) showing that, again, similarity in shape was the most predominant factor increasing intersection in their original positions.

### 4.6 Use of normalized CHV to study intersection dynamics in other neuronal groups

We use normalized CHPs to understand intersection dynamics in neurons that present dendritic domain size, shape, and cell body location that differ from our VTA DA neurons. We included mouse and rat SNc neurons, which we have described and show a similar architecture to VTA DA neurons (Henny et al., 2012; Montero et al., 2021), spinal motoneurons which are known to exhibit a very large and dense dendritic tree (Rotterman et al., 2014) and described as isodendritic (Ramón-Moliner & Nauta, 1966), and mediodorsal thalamic nucleus (Kuramoto et al., 2017), which exhibit much smaller dendritic domains. We show that intersection dynamics differ between cell types (Figure 7), for instance with a much less preponderant role of shape in MD and motoneurons. And we also show similar dynamics for both rat and mouse SNc DA neurons. We also include Purkinje cells (Nedelescu et al., 2018), which are characterized by extremely displaced cell bodies and a two-dimensionally arranged dendritic domain that generally does not overlap with neighboring neurons. In the context of this analysis, Purkinje cells work as a positive control for the method because, as expected from its sagittally organized two-dimensional dendritic domain, intersection in the ML axis falls rapidly.

### 4.7 Perspective

The broad spectrum of dendritic domain arrangements among neuronal types has been conceptualized as representing different types of connectivity, either of the selective, sampling, or space filling type (Harris & Spacek, 2017). The former type is characterized by small dendritic fields occupying a restricted locus, and represents an efficient solution to relay nervous signals from one neuronal stage to another as parallel, separate channels. The sampling strategy (and to some extent the space filling’s) connectivity strategy, on the other hand, is characterized by broader dendritic fields encompassing different loci, and represents a solution to increase integration and redundancy in signal transferring (Harris & Spacek, 2017). Our results allow us to predict that, for a given number of neurons and a given dendritic length, systems that require parallel and exclusive transferring of nervous signals may accomplish it by promoting heterogeneity in size, cell body location and specially shape of dendritic domains. Conversely, systems that require signal integration and redundancy may accomplish it by promoting uniformity in dendritic domain size, cell body location and shape.

## ACKNOWLEDGMENTS

We thank Luciana López-Juri for her insightful comments during the progress of this work. This work was supported by Fondo Nacional de Desarrollo Científico y Tecnológico Regular grants 1141170 and 1191497 to PH, and Comisión Nacional de Investigación Científica y Tecnológica REDES grant 180207 to PH.

## DATA AVAILABILITY STATMENT

Raw data, MATLAB scripts and data analysis are available at https://github.com/hennylab/Neuronal-Convex-hull-intersection and we have archived our code on Zenodo https://doi.org/10.5281/zenodo.6941258. SNc and VTA neurons reconstructions are available at https://www.neuromorpho.org.

## References

Arikkath, J. (2009). Regulation of dendrite and spine morphogenesis and plasticity by catenins. In Molecular Neurobiology (Vol. 40, Issue 1, pp. 46–54). Humana Press Inc. 10.1007/s12035-009-8068-x

Ascoli, G. A., & Krichmar, J. L. (2000). L-neuron: A modeling tool for the efficient generation and parsimonious description of dendritic morphology. Neurocomputing, 32–*33*, 1003–1011. 10.1016/S0925-2312(00)00272-1

Barber, C. B., Dobkin, D. P., & Huhdanpaa, H. (1996). The Quickhull Algorithm for Convex Hulls. ACM Transactions on Mathematical Software, 22(4), 469–483. 10.1145/235815.235821

Bodea, G. O., Spille, J. H., Abe, P., Andersson, A. S., Acker-Palmer, A., Stumm, R., Kubitscheck, U., & Blaess, S. (2014). Reelin and CXCL12 regulate distinct migratory behaviors during the development of the dopaminergic system. Development (Cambridge), 141(3), 661–673. 10.1242/dev.099937

Braitenberg, V., & Schüz, A. (1998). Cortex: Statistics and Geometry of Neuronal Connectivity. Springer Berlin Heidelberg. 10.1007/978-3-662-03733-1

Brignani, S., & Pasterkamp, R. J. (2017). Neuronal subset-specific migration and axonal wiring mechanisms in the developing midbrain dopamine system. In Frontiers in Neuroanatomy (Vol. 11). Frontiers Media S.A. 10.3389/fnana.2017.00055

Brombas, A., Kalita-De Croft, S., Cooper-Williams, E. J., & Williams, S. R. (2017). Dendro-dendritic cholinergic excitation controls dendritic spike initiation in retinal ganglion cells. Nature Communications, 8, 1–14. 10.1038/ncomms15683

Chazelle, B., & Dobkin, D. P. (1987). Intersection of convex objects in two and three dimensions. Journal of the ACM (JACM), 34(1), 1–27. 10.1145/7531.24036

Cuntz, H., Forstner, F., Borst, A., & Häusser, M. (2010). One rule to grow them all: A general theory of neuronal branching and its practical application. PLoS Computational Biology, 6(8). 10.1371/journal.pcbi.1000877

Cuntz, H., Mathy, A., & Häusser, M. (2012). A scaling law derived from optimal dendritic wiring. Proceedings of the National Academy of Sciences of the United States of America, 109(27), 11014–11018. 10.1073/pnas.1200430109

D’Errico, J. (2021). Inhull. https://www.mathworks.com/matlabcentral/fileexchange/10226-inhull

Fachada, N., & C. Rosa, A. (2018). micompm: A MATLAB/Octave toolbox for multivariate independent comparison of observations. Journal of Open Source Software, 3(23), 430. 10.21105/joss.00430

Farajian, R., Raven, M. A., Cusato, K., & Reese, B. E. (2004). Cellular positioning and dendritic field size of cholinergic amacrine cells are impervious to early ablation of neighboring cells in the mouse retina. Visual Neuroscience, 21(1), 13–22. 10.1017/S0952523804041021

Farassat, N., Costa, K. M., Stojanovic, S., Albert, S., Kovacheva, L., Shin, J., Egger, R., Somayaji, M., Duvarci, S., Schneider, G., & Roeper, J. (2019). In vivo functional diversity of midbrain dopamine neurons within identified axonal projections. ELife, 8. 10.7554/eLife.48408

Felix, R. A., Vonderschen, K., Berrebi, A. S., & Magnusson, A. K. (2013). Development of on-off spiking in superior paraolivary nucleus neurons of the mouse. Journal of Neurophysiology, 109(11), 2691–2704. 10.1152/jn.01041.2012

Fu, Y. H., Yuan, Y., Halliday, G., Rusznák, Z., Watson, C., & Paxinos, G. (2012). A cytoarchitectonic and chemoarchitectonic analysis of the dopamine cell groups in the substantia nigra, ventral tegmental area, and retrorubral field in the mouse. Brain Structure and Function, 217(2), 591–612. 10.1007/s00429-011-0349-2

Gärtner, U., Alpár, A., Reimann, F., Seeger, G., Heumann, R., & Arendt, T. (2004). Constitutive Ras activity induces hippocampal hypertrophy and remodeling of pyramidal neurons in synRas mice. Journal of Neuroscience Research, 77(5), 630–641. 10.1002/jnr.20194

Geisler, S., & Zahm, D. S. (2005). Afferents of the ventral tegmental area in the rat-anatomical substratum for integrative functions. Journal of Comparative Neurology, 490(3), 270–294. 10.1002/cne.20668

Gertler, T. S., Chan, C. S., & Surmeier, D. J. (2008). Dichotomous anatomical properties of adult striatal medium spiny neurons. Journal of Neuroscience, 28(43), 10814–10824. 10.1523/JNEUROSCI.2660-08.2008

Govender, N., Rajamani, R., Wilke, D. N., Wu, C. Y., Khinast, J., & Glasser, B. J. (2018). Effect of particle shape in grinding mills using a GPU based DEM code. Minerals Engineering, 129(September), 71–84. 10.1016/j.mineng.2018.09.019

Groves, P. M., & Linder, J. C. (1983). Dendro-dendritic synapses in substantia nigra: Descriptions based on analysis of serial sections. Experimental Brain Research, 49(2), 209–217. 10.1007/BF00238581

Harris, K. M., & Spacek, J. (2017). Chapter 1: Dendrite structure; in Dendrites (G. Stuart, N. Spruston, & M. Häusser, Eds.; 3th ed.). OUP Oxford.

Henny, P., Brown, M. T. C., Northrop, A., Faunes, M., Ungless, M. a, Magill, P. J., & Bolam, J. P. (2012). Structural correlates of heterogeneous in vivo activity of midbrain dopaminergic neurons. Nature Neuroscience, 15(4), 613–619. 10.1038/nn.3048

Herculano-Houzel, S. (2011). Not All Brains Are Made the Same: New Views on Brain Scaling in Evolution. Brain Behavior and Evolution, 78(1), 22–36. 10.1159/000327318

Hertel, N., Krishna-K, Nuernberger, M., & Redies, C. (2008). A cadherin-based code for the divisions of the mouse basal ganglia. Journal of Comparative Neurology, 508(4), 511–528. 10.1002/cne.21696

Hinds, J. W. (1970). Reciprocal and serial dendrodendritic synapses in the glomerular layer of the rat olfactory bulb. Brain Research, 17(3), 530–534. 10.1016/0006-8993(70)90263-5

Ito, T. (2020). Different coding strategy of sound information between GABAergic and glutamatergic neurons in the auditory midbrain. Journal of Physiology, 598(5), 1039–1072. 10.1113/JP279296

Jan, Y. N., & Jan, L. Y. (2010). Branching out: Mechanisms of dendritic arborization. Nature Reviews Neuroscience, 11(5), 316–328. 10.1038/nrn2836

Jones, B. (1995). Reticular Formation: Cytoarchitecture, Transmitters, and Projections. In G. Paxinos (Ed.), The Rat Nervous System (2nd ed., pp. 115–171). Academic Press.

Kawa, A., Stahlhut, M., Berezin, A., Bock, E., & Berezin, V. (1998). A simple procedure for morphometric analysis of processes and growth cones of neurons in culture using parameters derived from the contour and convex hull of the object. Journal of Neuroscience Methods, 79(1), 53–64. 10.1016/S0165-0270(97)00165-9

Kawano, H., Ohyama, K., Kawamura, K., & Nagatsu, I. (1995). Migration of dopaminergic neurons in the embryonic mesencephalon of mice. Developmental Brain Research, 86(1), 101–113. 10.1016/0165-3806(95)00018-9

Keeley, P. W., & Reese, B. E. (2010). Morphology of dopaminergic amacrine cells in the mouse retina: Independence from homotypic interactions. Journal of Comparative Neurology, 518(8), 1220–1231. 10.1002/cne.22270

Kline, S., & Felten, D. L. (1985). Ventral tegmental area of the rabbit brain: a developmental Golgi study. Brain Research Bulletin, 14(5), 485–492. 10.1016/0361-9230(85)90027-9

Kuramoto, E., Pan, S., Furuta, T., Tanaka, Y. R., Iwai, H., Yamanaka, A., Ohno, S., Kaneko, T., Goto, T., & Hioki, H. (2017). Individual mediodorsal thalamic neurons project to multiple areas of the rat prefrontal cortex: A single neuron-tracing study using virus vectors. Journal of Comparative Neurology, 525(1), 166–185. 10.1002/cne.24054

Lefebvre, J. L. (2017). Neuronal territory formation by the atypical cadherins and clustered protocadherins. In Seminars in Cell and Developmental Biology (Vol. 69, pp. 111–121). Elsevier Ltd. 10.1016/j.semcdb.2017.07.040

Lefebvre, J. L., Kostadinov, D., Chen, W. v., Maniatis, T., & Sanes, J. R. (2012). Protocadherins mediate dendritic self-avoidance in the mammalian nervous system. Nature, 488(7412), 517–521. 10.1038/nature11305

Lefebvre, J. L., Sanes, J. R., & Kay, J. N. (2015). Development of Dendritic Form and Function. Annual Review of Cell and Developmental Biology, 31, 741–777. 10.1146/annurev-cellbio-100913-013020

Malmierca, M. S., Seip, K. L., & Osen, K. K. (1995). Morphological classification and identification of neurons in the inferior colliculus: a multivariate analysis. Anatomy and Embryology, 191(4), 343–350. 10.1007/BF00534687

Meza, R. C., López-Jury, L., Canavier, C. C., & Henny, P. (2018). Role of the axon initial segment in the control of spontaneous frequency of nigral dopaminergic neurons in vivo. Journal of Neuroscience, 38(3), 733–744. 10.1523/JNEUROSCI.1432-17.2017

Michael Kleder. (2022). Centroid of a Convex n-Dimensional Polyhedron. https://www.mathworks.com/matlabcentral/fileexchange/8514-centroid-of-a-convex-n-dimensional-polyhedron

Mihaljević, B., Benavides-Piccione, R., Bielza, C., DeFelipe, J., & Larrañaga, P. (2015). Bayesian Network Classifiers for Categorizing Cortical GABAergic Interneurons. Neuroinformatics, 13(2), 193–208. 10.1007/s12021-014-9254-1

Montero, T., Gatica, R. I., Farassat, N., Meza, R., González-Cabrera, C., Roeper, J., & Henny, P. (2021). Dendritic Architecture Predicts in vivo Firing Pattern in Mouse Ventral Tegmental Area and Substantia Nigra Dopaminergic Neurons. Frontiers in Neural Circuits, 15. 10.3389/fncir.2021.769342

Muller, D. E., & Preparata, F. P. (1978). Finding the intersection of two convex polyhedra. Theoretical Computer Science, 7(2), 217–236. 10.1016/0304-3975(78)90051-8

Nedelescu, H., Abdelhack, M., & Pritchard, A. T. (2018). Regional differences in Purkinje cell morphology in the cerebellar vermis of male mice. Journal of Neuroscience Research, 96(9), 1476–1489. 10.1002/jnr.24206

Nieuwenhuys, R., Donkelaar, H. J., & Nicholson, C. (1998). The Central Nervous System of Vertebrates: With Posters (Issue v. 1-3). Springer.

Oleson, E. B., Cachope, R., Fitoussi, A., Tsutsui, K., Wu, S., Gallegos, J. A., & Cheer, J. F. (2014). Cannabinoid Receptor Activation Shifts Temporally Engendered Patterns of Dopamine Release. Neuropsychopharmacology, 39(6), 1441–1452. 10.1038/npp.2013.340

Packer, A. M., McConnell, D. J., Fino, E., & Yuste, R. (2013). Axo-dendritic overlap and laminar projection can explain interneuron connectivity to pyramidal cells. *Cerebral Cortex (New York*, N.Y.IZ: 1991), *23*(12), 2790–2802. 10.1093/cercor/bhs210

Peters, A., & Feldman, M. L. (1976). The projection of the lateral geniculate nucleus to area 17 of the rat cerebral cortex. I. General description. Journal of Neurocytology, 5(1), 63–84. 10.1007/BF01176183

Phillipson, O. T. (1979). A Golgi study of the ventral tegmental area of Tsai and interfascicular nucleus in the rat. Journal of Comparative Neurology, 187(1), 99–115. 10.1002/cne.901870107

Ramon y Cajal, S. (1904). Textura del sistema nervioso del hombre y de los vertebrados: estudios sobre el plan estructural y composición histológica de los centros nerviosos adicionados de consideraciones fisiológicas fundadas en los nuevos descubrimientos (Issue v. 2,n.°2). N. Moya.

Ramón-Moliner, E., & Nauta, W. J. H. (1966). The isodendritic core of the brain stem. Journal of Comparative Neurology, 126(3), 311–335. 10.1002/cne.901260301

Rice, M. E., & Patel, J. C. (2015). Somatodendritic dopamine release: Recent mechanistic insights. Philosophical Transactions of the Royal Society B: Biological Sciences, 370(1672), 1–14. 10.1098/rstb.2014.0185

Rojo, C., Leguey, I., Kastanauskaite, A., Bielza, C., Larrañaga, P., Defelipe, J., & Benavides-Piccione, R. (2016). Laminar Differences in Dendritic Structure of Pyramidal Neurons in the Juvenile Rat Somatosensory Cortex. Cerebral Cortex, 26(6), 2811–2822. 10.1093/cercor/bhv316

Rotterman, T. M., Nardelli, P., Cope, T. C., & Alvarez, F. J. (2014). Normal distribution of VGLUT1 synapses on spinal motoneuron dendrites and their reorganization after nerve injury. Journal of Neuroscience, 34(10), 3475–3492. 10.1523/JNEUROSCI.4768-13.2014

Takeo, Y. H., Shuster, S. A., Jiang, L., Hu, M. C., Luginbuhl, D. J., Rülicke, T., Contreras, X., Hippenmeyer, S., Wagner, M. J., Ganguli, S., & Luo, L. (2021). GluD2- and Cbln1-mediated competitive interactions shape the dendritic arbors of cerebellar Purkinje cells. Neuron, 109(4), 629–644.e8. 10.1016/j.neuron.2020.11.028

Tang, M., Miyamoto, Y., & Huang, E. J. (2009). Multiple roles of β-catenin in controlling the neurogenic niche for midbrain dopamine neurons. Development, 136(12), 2027–2038. 10.1242/dev.034330

Vrieler, N., Loyola, S., Yarden-Rabinowitz, Y., Hoogendorp, J., Medvedev, N., Hoogland, T. M., de Zeeuw, C. I., de Schutter, E., Yarom, Y., Negrello, M., Torben-Nielsen, B., & Uusisaari, M. Y. (2019). Variability and directionality of inferior olive neuron dendrites revealed by detailed 3D characterization of an extensive morphological library. Brain Structure and Function, 224, 1677–1695. 10.1007/s00429-019-01859-z

Zhang, L., Song, N.-N., Chen, J.-Y., Huang, Y., Li, H., & Ding, Y.-Q. (2012). Satb2 Is Required for Dendritic Arborization and Soma Spacing in Mouse Cerebral Cortex. Cerebral Cortex, 22(7), 1510–1519. 10.1093/cercor/bhr215

