## Supplementary Figures for "Geometrical factors determining dendritic domain intersection between neurons: a modeling study"

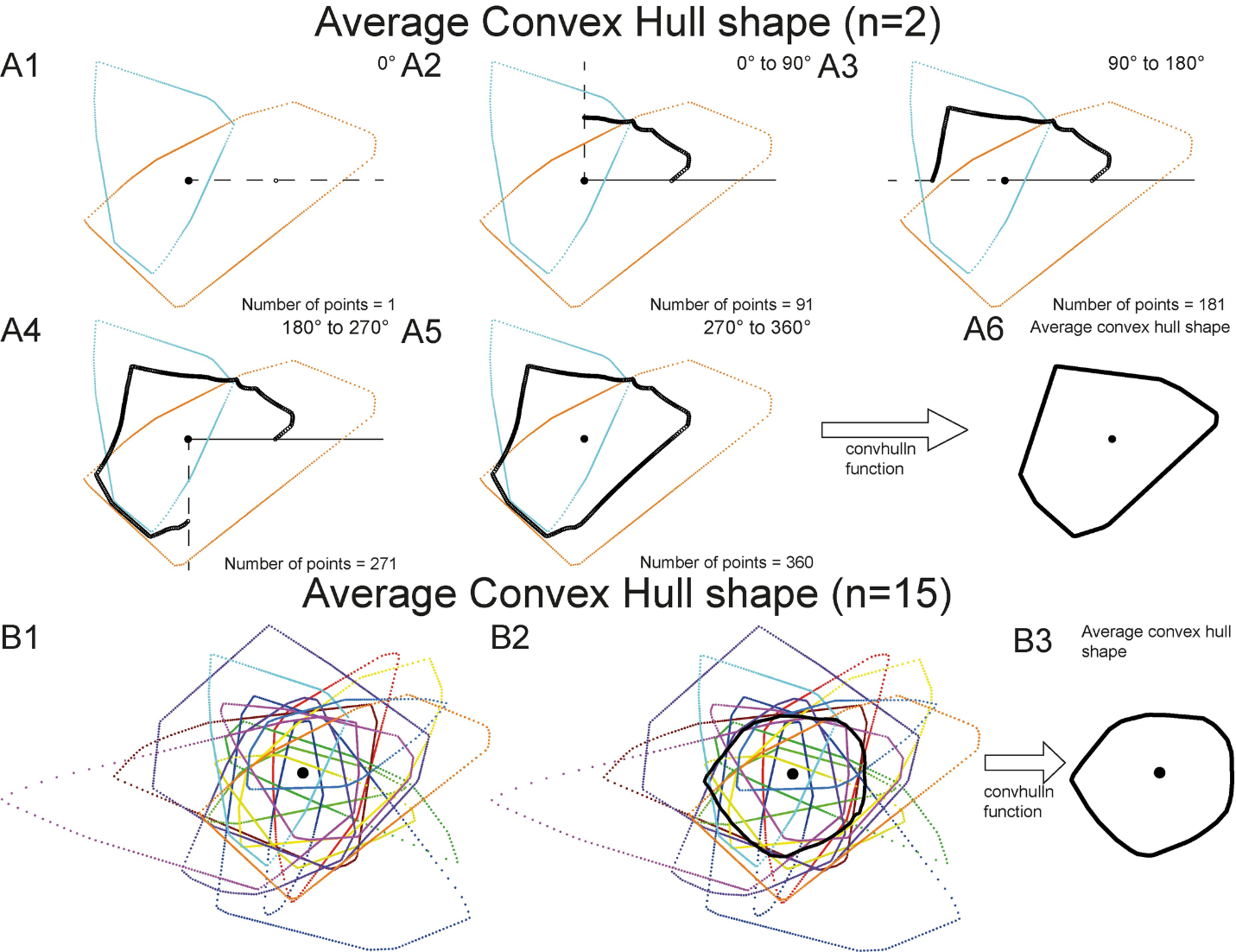

**Supplementary Figure 1. Simplified 2D representation of normalized shape computation.** A) Average shape method explained for a pair of 2D CH polygons. The polygons were centered in the origin in relation to their cell body, and a line is plotted at 0 degrees. The average point from the sample of 2 points in the given angle was obtained (black open circle, A1). Next, the process was repeated at 1-degree steps (A2-A5). Finally, 360 points were obtained, that were transformed to the average CHP shape using the *convhulln* function (A6). B) Average shape method explained for a group of 15 2D polygons. CHPs were centered in the origin (B1) and the average of the points in each angle were obtained (B2). Then, the average CHP shape was calculated using the *convhulln* function (B3). Scaling is different for examples B and C.

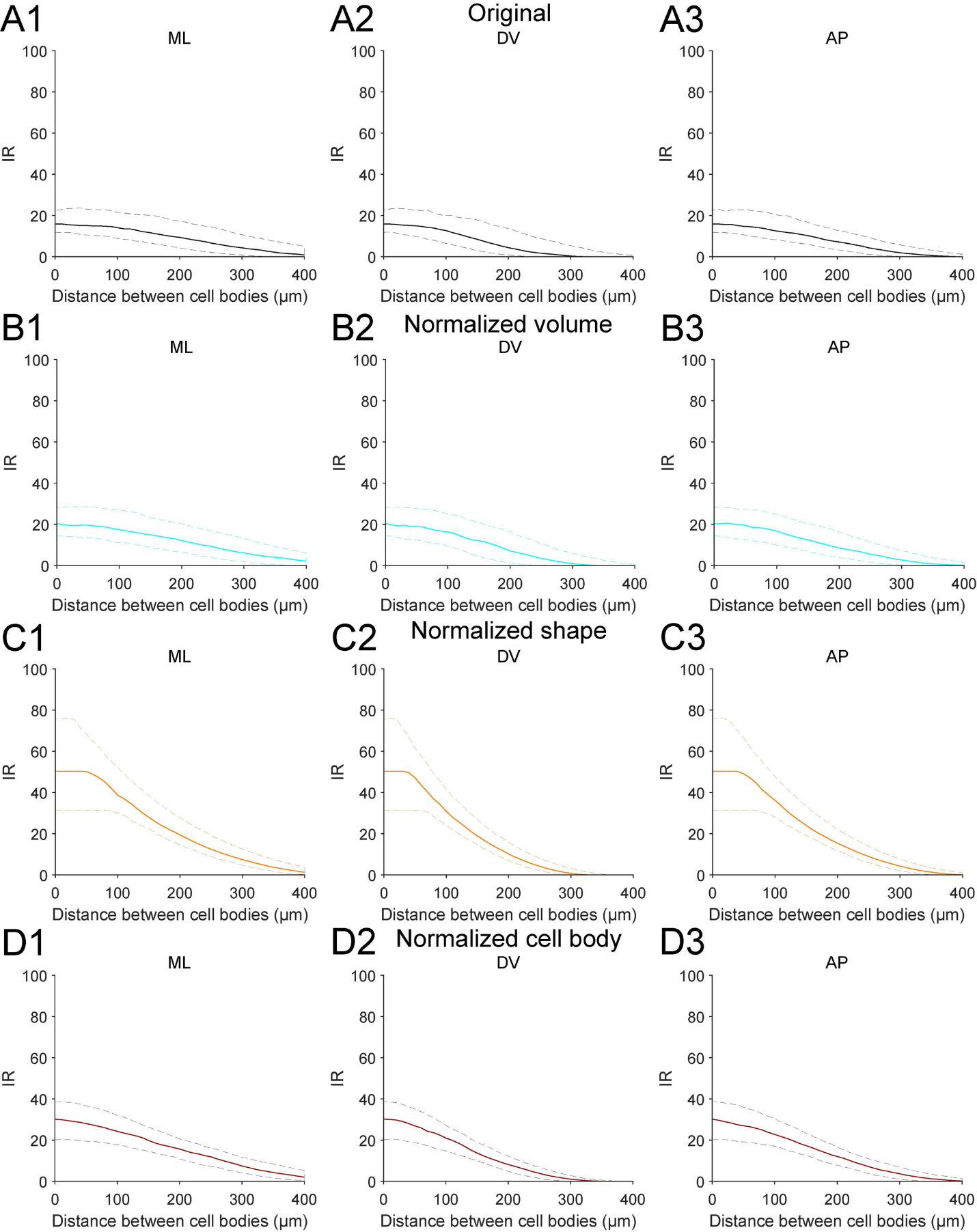

**Supplementary Figure 2. CHP modeling for VTA DA neurons: data with error bars.** IR vs distance between cell bodies plots across the ML , DV and AP axis were performed for all the possible pairs of VTA DA neurons (210 pairs per distance) for original data (A), normalized volume (B), normalized shape (C) and normalized cell body (D). IR was measured between 0 to 400 µm (10 µm steps). Solid line represents the median. Upper and lower dashed lines represent the 75th and 25th percentile, respectively.
